# Susceptibility monitoring and the molecular characterization of resistance of *Spodoptera frugiperda* (Lepidoptera: Noctuidae) to lambda-cyhalothrin and chlorpyrifos

**DOI:** 10.1101/2021.11.17.469006

**Authors:** Antonio Rogério Bezerra do Nascimento, Juliana Gonzales Rodrigues, Rubens H. Kanno, Fernando S.A. Amaral, José Bruno Malaquias, Karina Lucas Silva-Brandão, Fernando Luís Cônsoli, Celso Omoto

## Abstract

*Spodoptera frugiperda* (J. E. Smith) is a serious and widespread agricultural pest with several records of resistance to different insecticides and Bt proteins, including the neurotoxic insecticides chlorpyrifos (organophosphate) and lambda-cyhalothrin (pyrethroid). In this study, we (i) characterized and monitored the susceptibility of field populations of *S. frugiperda* to chlorpyrifos (194 populations) and lambda-cyhalothrin (197 populations) collected from major maize-growing regions of Brazil from 2003 to 2016, and (ii) compared gene expression levels of chlorpyrifos- and lambda-cyhalothrin-resistant strains to a susceptible reference strain (*Sf-ss*) of *S. frugiperda*. Laboratory-guided assays to monitor larval susceptibility detected average survival ranging from 29.3% to 36.0% to chlorpyrifos, and 23.1% to 68.0% to lambda-cyhalothrin at diagnostic concentration, based on LC_99_ of the susceptible reference strain of each insecticide. The resistance ratio of the chlorpyrifos-resistant strain (*Clo-rr*) was 25.4-fold and of the lambda-cyhalothrin-resistant strain (*Lam-rr*) was 217-fold. Differential gene expression analyses between resistant vs susceptible strains identified 1,098 differentially expressed genes (DEGs) between *Clo-rr* and *Sf-ss*, and 303 DEGs between *Lam-rr* and *Sf-ss*. Functional analyses of the DEGs revealed the up-regulation of several detoxification enzymes, mainly cytochrome P450 belonging to the *CYP3* and *CYP6* clans. Genes associated with regulatory processes, such as the forkhead box O (FoxO) were also up-regulated. Our data points that the resistance mechanisms of *Clo-rr* and *Lam-rr* strains of *S. frugiperda* to chlorpyrifos and lambda-cyhalothrin are mainly mediated by enzyme detoxification.

## Introduction

The fall armyworm (FAW) *Spodoptera frugiperda* (J.E. Smith) (Lepidoptera: Noctuidae) is native from the American continent but has recently turned into a global threat to agriculture after invading and spreading through the Old World (Baloch et al., 2020; CABI, 2021). The FAW is highly polyphagous feeding on more than 350 plant species (Montezano et al., 2018), and a voracious pest of several cash crops in the Americas, particularly maize (Montezano et al., 2018). There are several group of insecticides with different mode of action (carbamates, organophosphates, pyrethroids, neonicotinoids, spinosyns, benzoylureas, and diamides) and genetically modified crops expressing single or multiple toxins derived from the bacterium *Bacillus thuringiensis* (Bt-crops) (Cry1F, Cry1Ac, Cry1Ab, Cry2Ab2, Cry1A105 and Vip3a) available for FAW control in Brazil (AGROFIT, 2021). However, most of these technologies are at risk as there are reports of field-evolved resistance of S. frugiperda to Cry1F and Cry1Ab in Brazil (Farias et al., 2014; Omoto et al., 2016) and the presence of resistance alleles in natural populations that allowed the selection of resistant strains to different insecticides under laboratory conditions (Carvalho et al., 2013; Farias et al., 2014; Nascimento et al., 2016; Okuma et al., 2018; Bolzan et al., 2019; Lira et al., 2020; do Nascimento et al., 2021). The high capacity of this species to evolve resistance to several chemistries and different technologies of pest control requires a better understanding on the mechanisms of resistance involved for the appropriate use of different control strategies and the implementation of resistance management programs.

Neurotoxic insecticides have been widely used to control agricultural and urban pests. Pyrethroids are a large class of synthetic neurotoxic insecticides analog to pyrethrin, a substance present in daisy flowers, *Tanacetum cinerariifolium* (Matsuo, 2019). Pyrethroids inhibit the closure of voltage-gated sodium channels, maintaining the membrane of nervous cells in a continuous state of depolarization, resulting in paralysis of the organism (Soderlund and Bloomquist, 1989; Narahashi, 1996; Soderlund, 2005). Pyrethroids also induce autophagy and apoptosis in nerve cells (Park et al., 2017). Organophosphates (OP), a second group of neurotoxic insecticides acts by inhibiting acetylcholinesterase (AChE), an enzyme that catalyzes the hydrolysis of the neurotransmitter acetylcholine (ACh) at the synaptical cleft (Fukuto, 1990). Consequently, exposure to OP results in hyperexcitation of the nervous system (Spencer and O’Brien, 1957).

Prior to the advent of Bt crops, fall armyworm control was based on intensive use application of chemical insecticides. Unfortunately, the indiscriminate application of insecticides, mainly pyrethroids and organophosphates contributed to the evolution of *S. frugiperda* resistance to several compounds (Diez-Rodríguez and Omoto, 2001; Carvalho et al., 2013). High levels of resistance have been reported to the pyrethroids lambda-cyhalothrin, permethrin, cyhalothrin, tralomethrin, bifenthrin and fluvalinate (Diez-Rodríguez and Omoto, 2001; Carvalho et al., 2013), and to the organophosphates malathion, chlorpyrifos, methyl parathion, diazinon and sulprofos (Yu, 1991, 1992; Garlet et al., 2021). Resistance to pyrethroids and organophosphates has been commonly associated with mutations in genes coding for proteins of target sites and/or with modifications in the expression profile of genes encoding for detoxification enzymes, particularly monooxygenases of the cytochrome P450, esterases and glutathione-S-transferases (Scott, 1999; Daborn et al., 2002; Kasai, 2004; Djouaka et al., 2008; Feyereisen, 2012).

In this study, we monitored the susceptibility to the pyrethroid lambda-cyhalothrin and to the organophosphate chlorpyrifos in field populations of *S. frugiperda* collected from major maize-growing regions of Brazil from 2003 to 2016 and characterized the resistance of FAW to these neurotoxins. We applied large-scale transcriptome sequencing approach and identified the differentially expressed genes (DEGs) between lambda cyhalothrin- and chlorpyrifos-resistant strains and a susceptible reference strain for the characterization of the molecular mechanisms involved in FAW resistance to both neurotoxins. Detoxification of insecticides is an important adaptation of resistant strains to insecticides and toxins (Gordon, 1961; Li et al., 2007; Dermauw et al., 2013), and for this reason we evaluated the gene-expression profile of four main groups of detoxification enzymes related to insect resistance to pesticides: cytochrome P450s, glutathione-S-transferases, UDP-glycosyltransferases and esterases.

## Materials and Methods

### Susceptibility monitoring

We evaluated 194 and 197 field populations collected from major maize-growing regions from 2003 to 2016 to establish the larval susceptibility of *S. frugiperda* to chlorpyrifos and lambda-cyhalothrin, respectively (Supplementary Table 1). The diagnostic concentration used to monitor FAW susceptibility was 1000 μg ai/mL for chlorpyrifos and 56 μg ai/mL for lambda-cyhalothrin, based on LC_99_ of the susceptible reference strain of each insecticide (Carvalho et al. 2013). The field-collected insects were reared under laboratory conditions for one generation and the offspring obtained was subjected to bioassays. Bioassay included the topic application of 1 μL of the diagnostic concentration diluted in acetone onto the pronotum of fourth instars. Approximately 480 larvae were tested per population per insecticide (=40 replications of 12 larvae/population/insecticide). The bioassays were performed in 12-well acrylic plates (Costar® Corning Inc., Corning, NY, USA) containing artificial diet (Kasten et al., 1978). All plates were kept in a climate-controlled chamber at 25 ± 1 °C, 60 ± 10% RH and 14:10 h (L: D). The mortality was assessed after 24 h. Larvae were considered dead when not responding with coordinated movements when touched with a fine brush. A binomial generalized linear mixed model was used to compare the survival rate among seasons. We used the function *bootMer* from the package *lme4* (Bates et al., 2015)in the R package v.4.0.0 (R Core Team, 2020) to perform a mixed model-based semiparametric bootstrap and estimate the confidence intervals (95%) to each product.

### FAW susceptible and resistant strains

The susceptible reference strain of *S. frugiperda* (*Sf-ss*) has been maintained at the Arthropod Resistance Laboratory, Department of Entomology and Acarology, Luiz de Queiroz College of Agriculture (ESALQ/USP), Piracicaba, state of Sã o Paulo, Brazil without selection pressure from insecticides for more than 12 years. Larvae have been reared on an artificial diet based on beans, wheat germ, soy protein, yeast and casein (Kasten et al., 1978), and maintained in climate-controlled chambers (25 ± 1 °C; 60 ± 10% RH; 14:10 h L:D).

FAW strains resistant to lambda-cyhalothrin and chlorpyrifos were selected from insects collected in maize fields from the states of Paraná (PR) and Bahia (BA), respectively, by using the F_2_ screening method (Andow and Alstad, 1998). A total of 72 (PR) and 86 (BA) two-parent isofamilies were established from the field-collected insects. Fourth instars of these isofamilies were used for the application of 1 μL of an acetone solution of chlorpyrifos (1000 μg ai/mL) (99% pure, Dow AgroSciences) or lambda-cyhalothrin (56 μg ai/mL) (87.4% pure, Syngenta) on their pronotum using a micro-applicator. Larvae were transferred to acrylic plates and maintained under laboratory-controlled conditions as earlier described. After 24 h, the surviving FAW larvae were transferred to plastic cups containing an artificial diet (Kasten et al., 1978) to complete their life cycle. Insects that survived insecticide exposure at the tested concentrations were considered the parental resistant strain.

### Characterization of *S. frugiperda* chlorpyrifos- and lambda-cyhalothrin-resistant strains

The toxicological characterization of the *Sf-ss*, *Clo-rr* and *Lam-rr* strains was performed using 8 to 12 logarithmically spaced concentrations of each insecticide as earlier described. Control larvae were treated only with acetone. The experimental design was completely randomized, with five replicates (12 larvae/replicate) totaling 60 larvae tested per concentration. Mortality was assessed after 24 h of insecticide application, and subjected to Probit analysis (Finney, 1971) using the software POLO PLUS (LeOra Software, 1987). The resistance ratio was estimated by dividing the LC_50_ of the resistant strain by the LC_50_ of the *Sf-ss* strain. Tests for parallelism and equality of regression constants were also conducted.

### Total RNA extraction, library preparation and Illumina sequencing

Total RNA was isolated from three replicates of four larvae (=50 mg)/each of the *Sf-ss*, *Lam-rr* and *Clo-rr* strains. Larvae were collected in 700 μL of Trizol^™^ Reagent (Invitrogen^®^, Carlsbad, CA, USA), subjected to mechanical maceration and centrifugation (16,000 *g* × 5 min). The supernatant was collected and added to 700 μL of 95% ethanol for total RNA recovery using the Direct-zol^™^ RNA mini-prep kit (Zymo Research^®^, Irvine, CA, USA). The solution was transferred to filter columns and centrifuged (16,000 *g* × 30 s). Subsequently, 400 μL of RNA wash buffer, 5 μL of DNAse I (6 U.μL^−1^) and 75 μL of DNA digestion buffer were added to the membrane, followed by incubation at room temperature for 15 min. Afterwards, 400 μL of Direct-zol^™^ RNA PreWash (Zymo Research^®^, Irvine, CA, USA) was added and the column was centrifuged (16,000 *g* × 30 s). Finally, 700 μL of Wash Buffer RNA was added and the tubes were centrifuged at 16,000 *g* for 2 min or until the wash buffer was completely removed. RNA was eluted in 50 μL of DNA/RNA-free water and stored at –80 °C until further use.

Samples were sent to the Central Laboratory of High-Performance Technologies in Life Sciences (LACTAD/UNICAMP) for evaluation of RNA quality and integrity, library preparation and sequencing in an Illumina HiSeq2500^®^ platform using the paired-end strategy (2 × 100 bp).

### Reads quality-filtering, mapping and annotation

Reads were quality-assessed using FastQC v0.11.5 (Andrews, 2010). Illumina adapter sequences and low-quality reads (Phred quality score < 20) were trimmed with Trimmomatic v 0.36 (Bolger et al., 2014). The transcriptome was assembled by mapping the trimmed reads against the draft genome of *S. frugiperda* as a reference (Gouin et al., 2017) using Bowtie v.2 (Langmead and Salzberg, 2012). To maximize the functional identification, the assembled transcripts were annotated using BLASTX search with Diamond (Buchfink et al., 2015) against the *non-redundant* (nr)-database of the NCBI. Enzyme classification (EC) codes and the annotation of metabolic pathways KEGG (Kyoto Encyclopedia of Genes and Genomes) (Kanehisa et al., 2007) were generated with Blast2GO (Conesa et al., 2005) with an *e*-value cut-off set to 10^−3^. The transcripts were blasted against the GO (Gene Ontology), EggNOG (Powell et al., 2011)and UniProt databases, with an *e*-value cut-off set to 10^−5^.

### Identification of DEGs

Differentially expressed genes (DEGs) between resistant and susceptible strains of *S. frugiperda* were determined based on expression abundancies in each condition. The calculation of relative abundance was obtained by counting reads mapped against the reference genome (https://bipaa.genouest.org/sp/spodoptera_frugiperda_pub/download/genome/Pseudo_genome/) using RSEM v 1.1.17 (Li and Dewey, 2011) to estimate the expression abundance of genes and isoforms by determining the number of fragments per kilobase of exon per million of mapped fragments (FPKM). The differential expression analysis was evaluated using the DESeq2 package (Love et al., 2014). The data for expression-abundance estimation were normalized using correction factors based on the effective size of the libraries.

## Results

### Susceptibility monitoring

The overall susceptibility of 197 field-populations of *S. frugiperda* to lambda-cyhalothrin increased over the seasons for the field populations tested, with the mean survival ranging from 23.1% in the 2003 season to 68.0% in 2016. The mean survival of 194 field populations tested with chlorpyrifos ranged from 29.3% in the 2003 to 36.0% in 2016 season (Figure 1A and 1B). In general, populations collected from the states of Bahia in areas within the limits of the Brazilian savanna (*Cerrado*) presented the lower susceptibility to both insecticides tested (Figure 1C and 1D).

**Figure 1.**
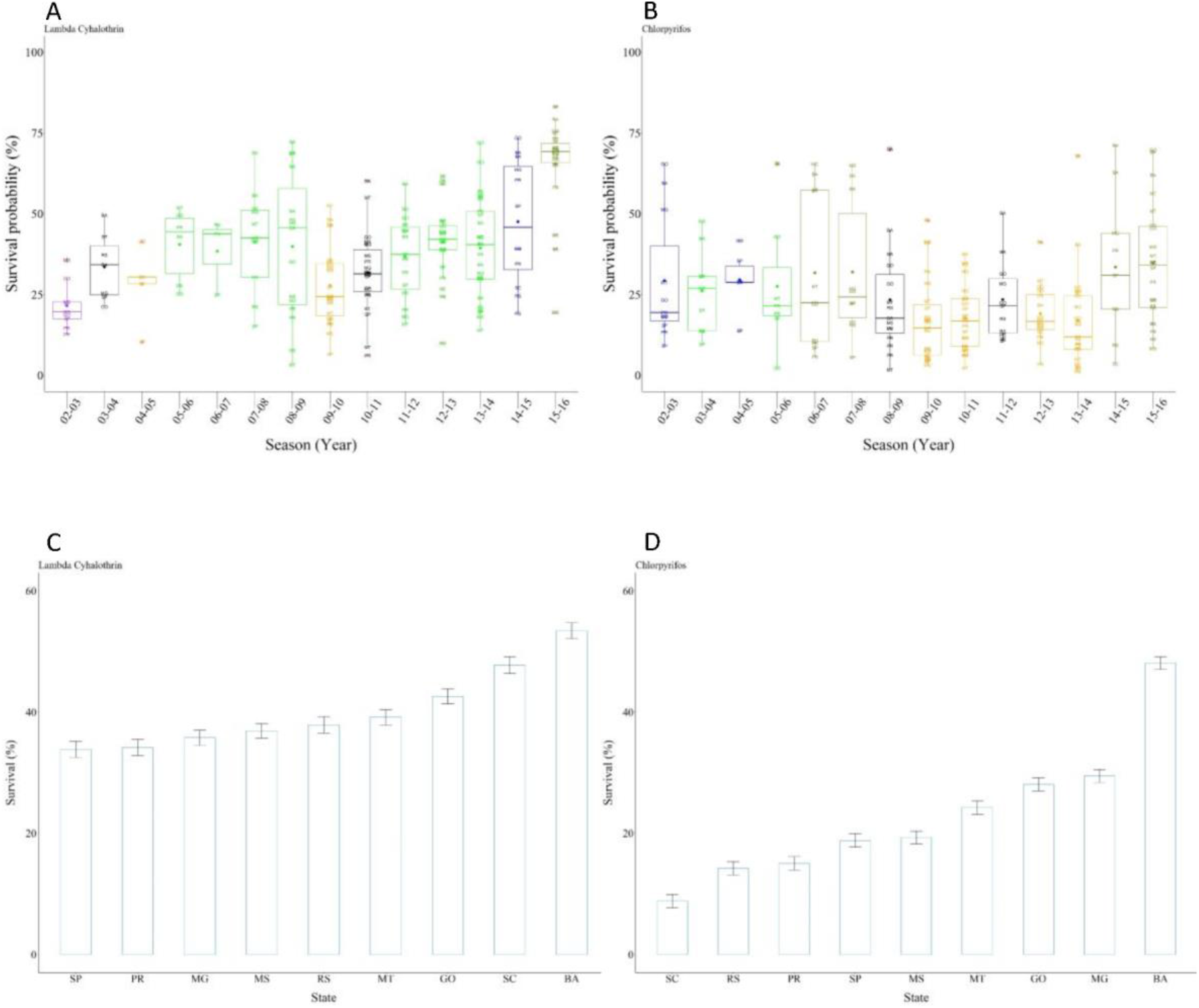
Monitoring of susceptibility of *S. frugiperda* to lambda-cyhalothrin (A) and chlorpyrifos (B). Boxplots with the same color (group) are not significantly different when the confidence interval overlaps (95% CI). The 95% CIs were estimated with a binomial generalized linear mixed model. Outliers are shown as asteristics (*) out of boxplots, while mean values are shown as filled circle in the boxplots. Mean of susceptibility of *S. frugiperda* to lambda-cyhalothrin (C) and chlorpyrifos (D) by the Brazilian States.

### Characterization of laboratory-selected lambda-cyhalothrin and chlorpyrifos resistant strains

Concentration-response bioassays of laboratory-selected *Clo-rr* and *Sf-ss* larvae of *S. frugiperda* to chlorpyrifos resulted in LC_50_ values of 854.41 μg.mL^−1^ (617.35 – 1,236.96) for *Clo-rr* and of 33.64 μg.mL^−1^ (8.70 – 71.36) for *Sf-ss*, with 25.4-fold resistance ratio. Similar dose-response assays of *Lam-RR* and *Sf-ss* with lambda-cyhalothrin resulted in LC_50_ values of 76.18 μg.mL^−1^ (49.34 – 107.21) for *Lam-rr* and only 0.35 μg.mL^−1^ (0.09 – 0.86) for *Sf-ss*, with 217.6-fold resistance ratio (Table 1).

**Table 1.** Concentration – mortality of susceptible and resistant strains of *S. frugiperda* to lambda-cyhalothrin and chlorpyrifos.

| Strain (insecticide) | n | Slope (±SE) | LC<sub>50</sub> (CI 95%) | $\chi^2$ | df | RR |
| --- | --- | --- | --- | --- | --- | --- |
| *Sf-ss* (Lambda-cyhalothrin) | 468 | 0.63 ± 0.07 | 0.35 (0.09 – 0.86) | 13.80 | 5 | - |
| *Lam-rr* (Lambda-cyhalothrin) | 504 | 2.92 ± 0.23 | 76.18 (49.34 – 107.21) | 8.78 | 3 | 217.6 |
| *Sf-ss* (Chlorpyrifos) | 488 | 1.81 ± 0.28 | 33.64 (8.70 – 71.36) | 5.29 | 7 | - |
| *Clo-rr* (Chlorpyrifos) | 742 | 2.05 ± 0.13 | 854.41 (617.35 – 1236.96) | 16.29 | 5 | 25.4 |

1. \*LC<sub>50</sub> values followed by the same letter in the columns do not differ significantly for the confidence intervals (95%). The significance of the confidence intervals was determined by the likelihood ratio test, followed by multiple comparisons. 2. \*\*df = degrees of freedom. 3. \*\*\*Resistance ratio (RR) = LC<sub>50</sub> of resistant strain/LC<sub>50</sub> of susceptible strain.

### Re-annotation of *S. frugiperda* genome

The re-annotation of genes from the draft genome increased the number of genes identified (Supplementary Figure 1). A total of 19,600 open reading frames (ORFs) were annotated using NR-NCBI; 11,902 in KOG; 6,801 in GO; and 6,381 in KEGG. The total number of ORFs annotated was 19,621, with the annotation of 8,579 additional ORFs than previously reported (Gouin et al., 2017).

### DEGs in lambda-cyhalothrin-resistant and susceptible strains

Comparative analysis between the transcriptomes of the *Lam-rr* and *Sf-ss* strains identified a total of 303 DEGs (Supplementary Table 2), out of which 176 were up-regulated and 127 down-regulated in *Lam-rr*. Annotation indicated 10% of DEGs was related to metabolism and carbohydrate transport, with the categories signal-transduction mechanism, energy production and conversion, and lipid transport and metabolism representing almost 5% of the transcripts (Figure 2A and B). Differential expression analysis identified the up-regulation of the subfamilies *CYP321B1*, CYP*6AE44*, *CY321A8*, *CYP321A10* and *CYP321A7* of the cytochrome P450 monooxygenase in *Lam-rr* larvae (log_2_ FC ranging from 2.01 to 3.61) (Fig. 3A). GSTs *epsilon* 2, *epsilon* 9 and *sigma* 2, the subfamily of the UDP-glycosyltransferase 39B4, and six esterase transcripts were up-regulated in *Lam-rr* (Figure 3B, C and D).

**Figure 2.**
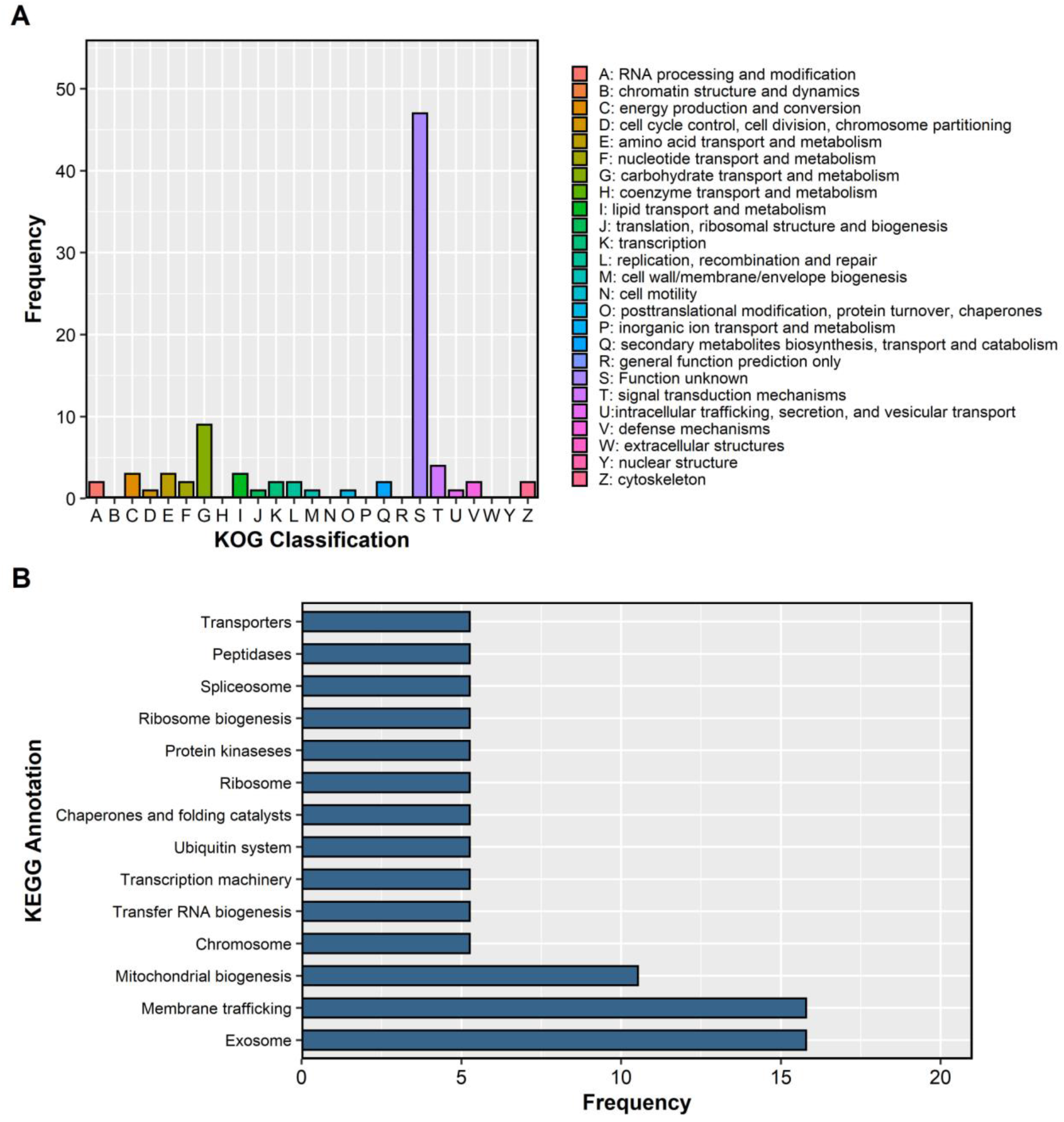
Distribution of DEGs between resistant and susceptible strains of *S. frugiperda* to lambda-cyhalothrin annotated by KOG databases (A) and KEGG (B)

**Figure 3.**
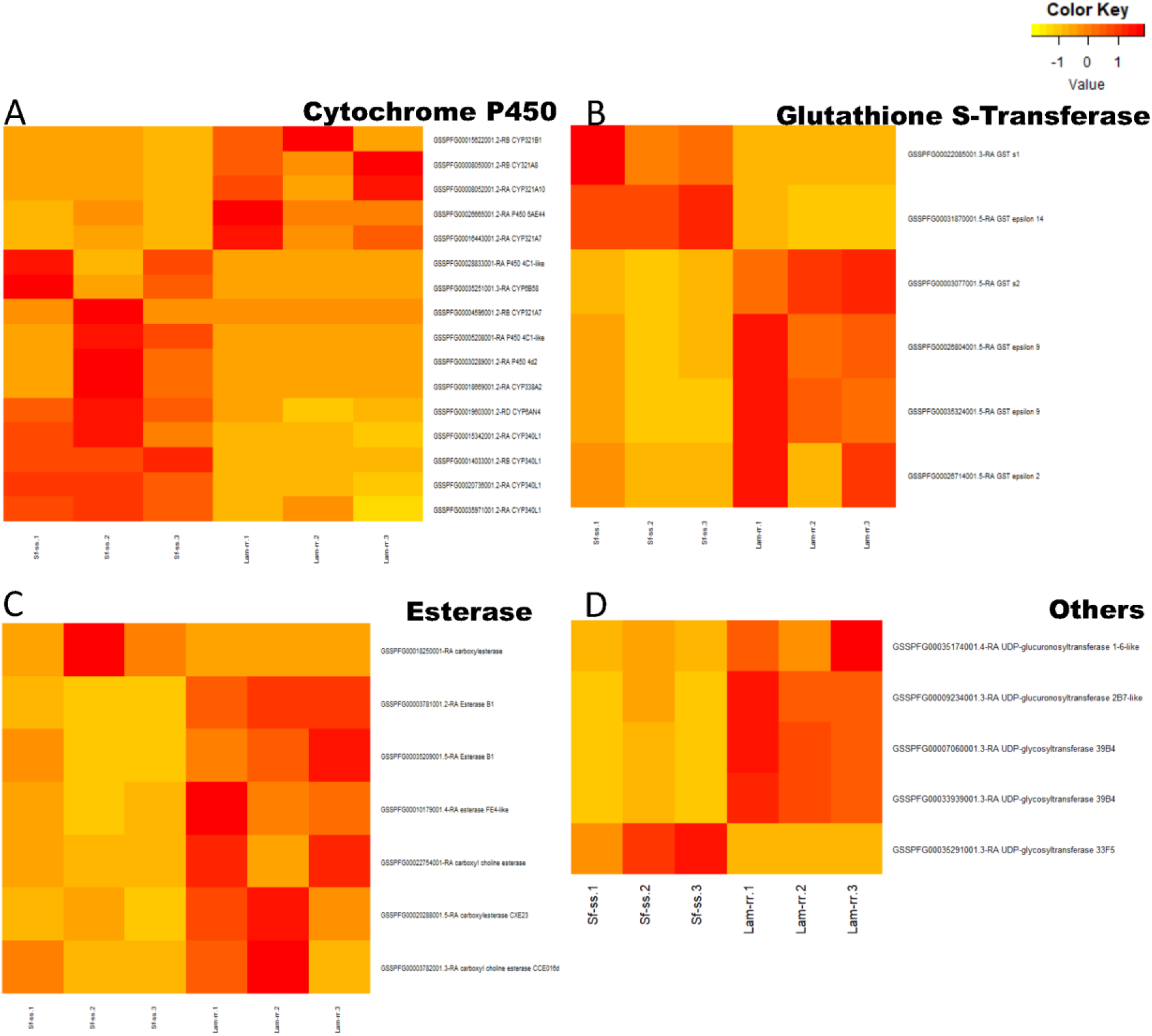
Heatmap of differentially expressed cytochrome P450s (A), glutathione S-transferases (B), esterases (C) and other detoxification enzymes (D) when comparing susceptible and resistant *S. frugiperda* strains to lambda-cyhalothrin

### DEGs between chlorpyrifos-resistant and susceptible strains

The comparative analysis between *Clo-rr* and *Sf-ss* strains identified 1,098 differentially expressed genes (DEGs), with the up-regulation of 601 and the down-regulation of 497 in *Clo-rr* larvae (Supplementary Table 3). The functional annotation showed that amino acid, lipid, and carbohydrate transport were highly represented among the DEGs. Functions associated with regulatory processes with post-transcriptional and transduction signals were also frequent among DEGs (Figure 4A). KEGG pathway analysis showed that the metabolism and biosynthesis of secondary metabolites pathways were the most represented (Figure 4B).

**Figure 4.**
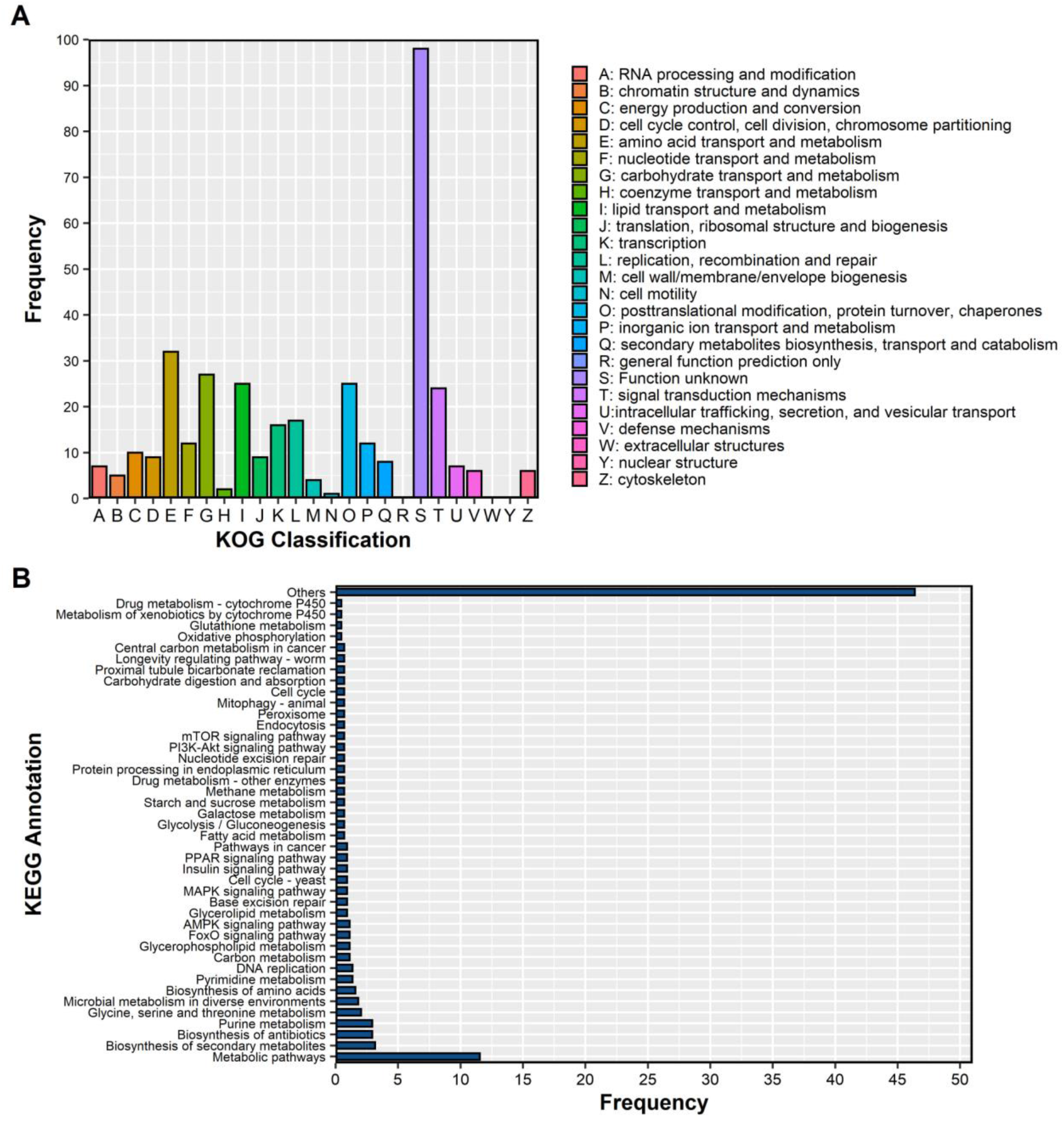
Distribution of DEGs between resistant and susceptible strains of *S. frugiperda* to chlorpyrifos annotated by KOG databases (A) and KEGG (B)

GO term enrichment showed the molecular function categories urate oxidase and oxidoreductase activities (GO:0016701, GO:0016702, GO:0016661, GO:0016663) were over-represented in DEGs of *Clo-rr*, while the biological processes and the cellular component PcG proteins complex were under-represented (Figure 5A, B and C). Genes related to oxidative stress response Strain pathway were also common among DEGs. Many key genes related to stress response were up-regulated in *Clo-rr S. frugiperda* larvae, such as G-protein coupled receptors (GPCR) (log_2_ FC = 2.38), cAMP-responsive element binding (CREB) (log_2_ FC = 2.84) and forkhead box `other` (FoxO) (log_2_ FC = 2.13) (Supplementary Table 3). Enrichment analysis also detected a set of transcripts of detoxification enzymes commonly associated with insecticide resistance in insects, which were up-regulated in the *Clo-rr* strain. These enzymes belong to subfamilies of cytochrome P450 monooxygenase (*CYP*), including *CYP333B3, CYP367A6, CYP340AA1, CYP6AB14, CYP49A1* and *CYP6A2* (Figure 6A), glutathione S-transferase epsilon 14 (GST) (Figure 6B), esterase (Figure 6C) and UDP-glycosyltransferase 39B4, 50A5 (UGT) (Figure 6D).

**Figure 5.**
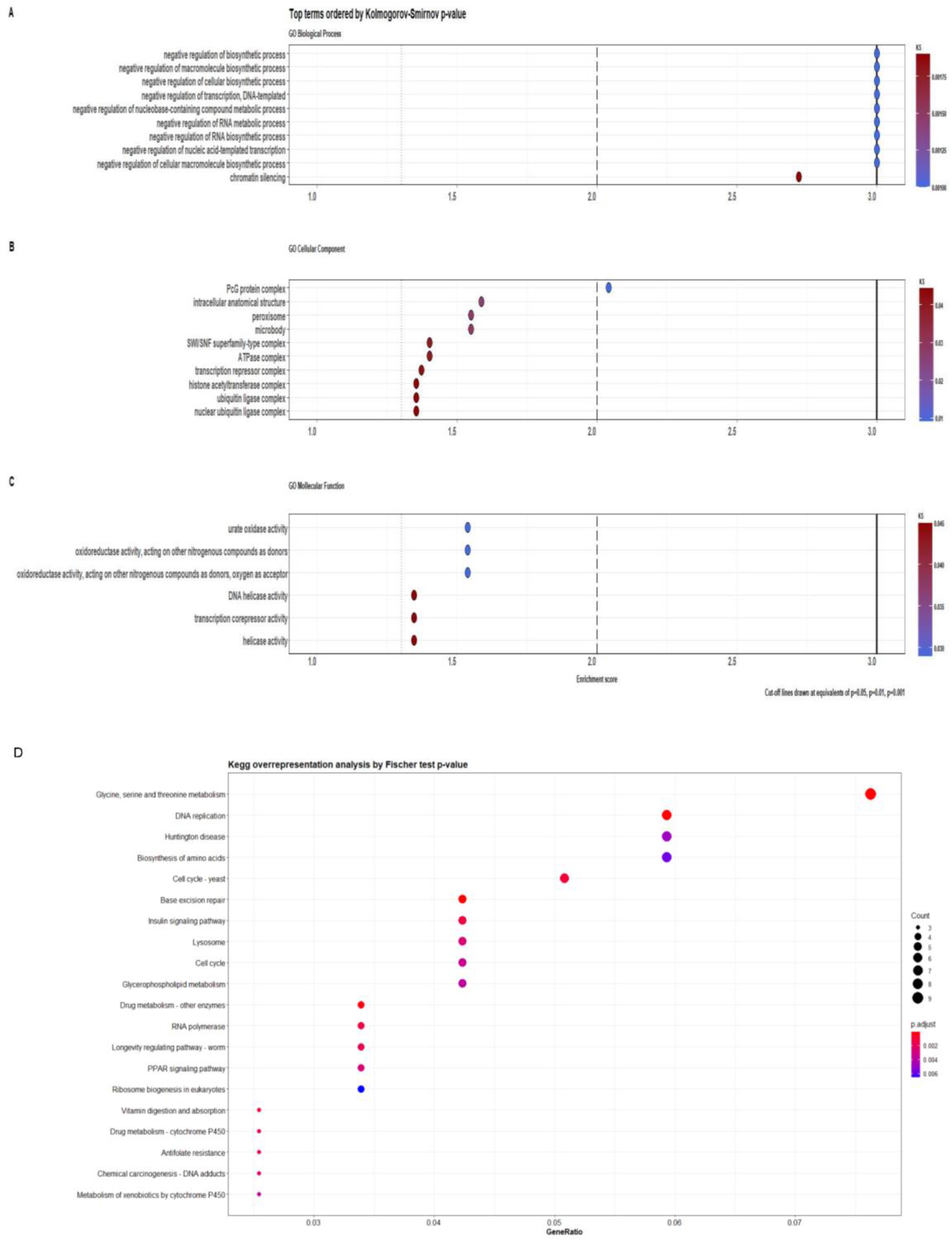
The 10 best enriched Gene Ontologies (GOs) for the DEGs between resistant and susceptible strains of *S. frugiperda* to chlorpyrifos (A, B and, C) and Kegg overrepresentation analysis (D).

**Figure 6.**
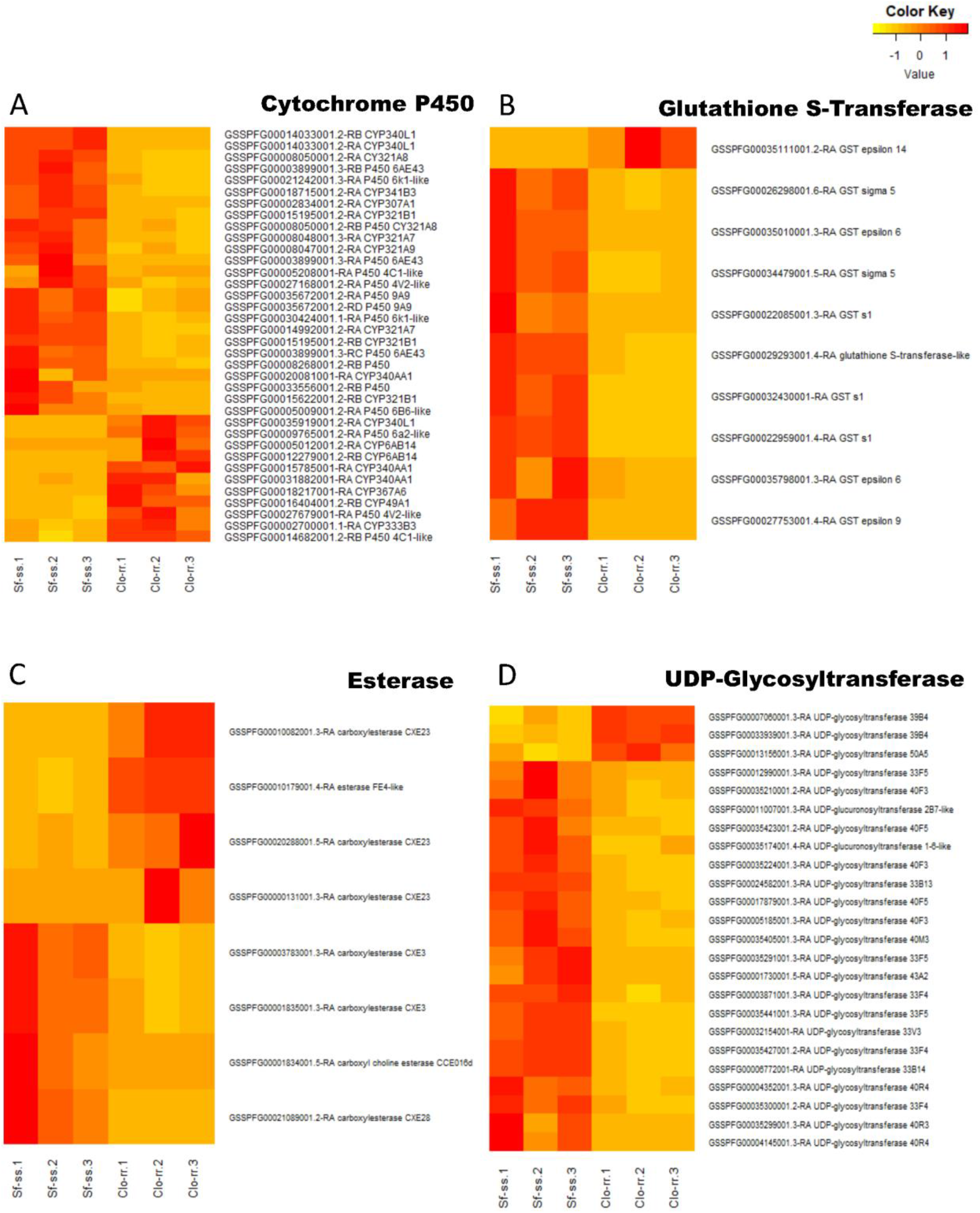
Heatmap of differentially expressed cytochrome P450s (A), Glutathione S-transferases (B), esterases (C) and UDP-glycosyltransferases (D) when comparing susceptible and resistant *S. frugiperda* strains to chlorpyrifos.

## Discussion

The susceptibility of field-collected populations of *S. frugiperda* to the neurotoxic insecticides lambda-cyhalothrin and chlorpyrifos was reduced from 2003 to 2016, allowing the selection of resistant strains and the investigation of the resistance mechanisms of *S. frugiperda* to both insecticides using RNA-seq analyses.

The strong selection pressure caused by the continuous use of insecticides resulted in the evolution of resistance of *S. frugiperda* to several control strategies, including the intensive and uninterrupted use of pyrethroids in pest control in Brazil. Pyrethroids represent a significant portion of the market share of pesticides in Brazil (Personal information, Spark Smarter Decisions-http://spark-ie.com.br/?lang=en), and it is often used in pest control in maize, soybean, and cotton. Maize is available in the field almost all year round, and maize fields overlap with those of soybean and cotton, as these crops are cultivated simultaneously or successively, particularly in the Brazilian savanna. Moreover, pyrethroids are also recommended to control other insect pests such as stink bugs, whiteflies, and leafhoppers besides lepidopteran pests (AGROFIT, 2021). These factors contributed altogether for the high selection pressure with insecticides favoring the evolution of resistance of *S. frugiperda* throughout growing seasons.

The existence of multiple factors involved with the evolution of resistance of *S. frugiperda* to lambda-cyhalothrin is not as clear as that to the organophosphate chlorpyrifos. Monitoring *S. frugiperda* survival frequency over 15 crop seasons demonstrated a stabilization of the susceptibility to chlorpyrifos. Chlorpyrifos is also a broadly used insecticide, having a significant share of the insecticides marketed in Brazil, and *S. frugiperda* is certainly under high selection pressure due to the continuous exposure to this pesticide (IBAMA, 2021)The continuous exposure of *S. frugiperda* to the selection pressure of this pesticide, and the steady low survival probability observed during our monitoring indicate that resistance to chlorpyrifos is associated with very high fitness costs. In fact, analysis of the fitness costs of chlorpyrifos resistance in a laboratory-selected strain of *S. frugiperda,* demonstrated resistance affected several biological traits at the individual and population levels (Garlet et al., 2021). The fitness costs associated with chlorpyrifos resistance in *S. frugiperda* is so high that Garlet et al. (2021) concluded populations of *S. frugiperda* would restore their susceptibility to chlorpyrifos with the suspended use of the selecting agent. The fitness cost associated with chlorpyrifos resistance is also reported to several other insect species, and in chlorpyrifos-resistant *Plutella xylostella* (Lepiodptera: Plutellidae) it has been suggested to occur due the excessive energy use for the over-production of detoxification enzymes and heat shock proteins when these proteins are not required (Zhang et al., 2015; Yang et al., 2018).

A large set of DEGs identified in comparative analyses of chlorpyrifos- and lambda-cyhalothrin-resistant strains with their respective susceptible strains is involved with the first two steps of the process of detoxification. Several subfamilies of *CYP3* and *CYP6* clades were up-regulated in chlorpyrifos- and lambda-cyhalothrin-resistant strains. Monooxygenases of the cytochrome P450 are often involved in xenobiotic detoxification in insects, including a large set of different organic insecticide molecules (Feyereisen, 1999; Lu et al., 2021; Nauen et al., 2021). Differential gene expression analyses through massive sequencing of cDNA libraries of insecticide-resistant strains often reveals a large number of *CYP* genes that are overexpressed in resistant as compared to susceptible insects (Djouaka et al., 2008; Brun-Barale et al., 2010; Nascimento et al., 2015). Some studies have confirmed the role of CYP enzymes in the detoxification of insecticides, and consequently in the resistance mechanism of several species by using enzyme inhibitors (Rose et al., 1995; Scharf et al., 1997; Liu and Scott, 1998), genomic analysis and broad RT-qPCR experiments (Yang and Liu, 2011), and gene editing and/or gene knockout (Itokawa et al., 2016; Shi et al., 2018; Wang et al., 2018a).

The CYP clades most represented in our differential gene expression analysis have been directly involved in resistance to pyrethroids and organophosphates. *CYP3* genes were shown to have a high constitutive expression in the integument of deltamethrin-resistant *Triatoma infestans* (Hemiptera: Reduviidae), and to participate in the process of detoxification of this pyrethroid at the insect integument (Dulbecco et al., 2018). Knockdown of CYP6 genes were also shown to increase *Locusta migratoria* (Orthoptera: Acrididae) susceptibility to carbamates and pyrethroids (Zhang et al., 2019). Overexpression of multiple CYP genes were also involved in detoxification of organophosphates, including chlorpyrifos (Sabourault et al., 2001; Brun-Barale et al., 2010; Wang et al., 2018b; Bo et al., 2020). However, chlorpyrifos oxidation by a particular *CYP6* gene (CtCYP6EX3) in *Chironomus tentans* (Diptera: Chironomidae) was demonstrated to lead to an increased larval mortality due the formation of chlorpyrifos-oxon, a metabolite of increased toxicity to *C. tentans* (Tang et al., 2017). The participation of CtCYP6EX3 in the toxicity of chlorpyrifos to *C. tentans* was detected after the verification of the up-regulation of several *CYP* genes after insect exposure to the herbicide atrazine, with an antagonist effect on the efficiency of organophosphates (Tang et al. 2017).

Insect resistance to lambda-cyhalothrin and chlorpyrifos can also result from site mutations. There is a large number of point mutations in the voltage-sensitive sodium channel leading to target insensitivity to pyrethroids (Dong et al., 2014), including lambda-cyhalothrin (Wu et al., 2014; Zhen and Gao, 2016; Tian et al., 2018). Several point mutations were reported in the acetylcholinesterase 1 gene in chlorpyrifos-resistant insects, such as G119S in *Culex quinquefasciatus* (Diptera: Culicidae) (Liu et al., 2005), F392W in *Bemisia tabaci* (Hemiptera: Aleyrodidae) (Zhang et al., 2012), F439H in *Laodelphax striatellus* (Homoptera: Delphacidae) (Zhang et al., 2013), G119S and F331C in *Nilaparvata lugens* (Homoptera: Delphacidae) and A201S in *Tuta absoluta* (Lepidoptera: Gelechiidae) (Haddi et al., 2017). Resistance to organophosphates was also related to structural mutations in carboxylesterases, resulting in an enhanced kinetics for insecticide hydrolysis (Oakeshott et al., 2005).

Despite the long list of point mutations in the target sites of lambda-cyhalothrin and chlorpyrifos, the mechanism involved in resistance in the selected strains of *S. frugiperda* is not related to point mutations and target-site insensitivity. Contrarily to a previous report (Carvalho et al., 2013), we did not detect non-synonymous mutations in the sodium channel and acetylcholinesterase 1 of *S. frugiperda* strains resistant to lambda-cyhalothrin or chlorpyrifos, respectively. Moreover, the overproduction of detoxification enzymes as the underlying mechanism in lambda-cyhalothrin and chlorpyrifos resistance of *S. frugiperda* is also supported by the up-regulation of detoxification enzymes other than CYP. The over-expression of glutathione-S-transferases, esterases and UDP-glucuronosyltransferases potentiates the hydrolysis and modification of these pesticides in the process of detoxification of these pesticides (Samra et al., 2012). Additionally, GSTs can also play an important role in insect resistance by binding and passively sequestering insecticide molecules (Kostaropoulos et al., 2001; Kristensen, 2005; Sonoda and Tsumuki, 2005; Lumjuan et al., 2011; Hsu et al., 2016), protecting tissues by reducing the oxidative stress and lipid peroxidation induced by pyrethroids (Feyereisen, 1999; VONTAS et al., 2001).

But most differentially expressed GSTs and UGTs were down-regulated in the *Clo-rr* strain. Down-regulation of GSTs upon chlorpyrifos exposure was already reported in *L. migratoria*, with the up-regulation of a single GST gene (Qin et al., 2014). GSTs are classified into six major classes. Four (*sigma*, *omega*, *zeta* and *theta*) of them are common to metazoans, while the remaining two (*delta* and *epsilon*) are specific to insects (Friedman, 2011). Changes in gene expression and/or activity of the *delta* and *epsilon* classes of GSTs are the most commonly associated with insecticide resistance in insects (Enayati et al., 2005; Lumjuan et al., 2011; Pavlidi et al., 2018), which would explain the increased gene expression observed in *Clo-rr*, since the only up-regulated GST was the *epsilon* 14. A similar pattern of regulation of UGTs genes was observed for *Clo-rr*. Changes in UGTs expression were detected in 14 subfamilies, but only *UGT39B4* was up-regulated in *Clo-rr*. Gene expression of UGTs in *S. exigua* larvae treated with different insecticides varied with the type of insecticide used, with 46.9 to 84.4% of the 32 UGT genes tested showing no alteration in expression after insect exposure, 3.1 to 46.9% were down-regulated and 6.2 to 21.9% were up-regulated. Tissue expression analysis indicated *UGT39B4* was overexpressed in the head when compared to midgut, Malpighian tubules and fat body (Hu et al., 2019).

Moreover, the up-regulation of carboxylesterases (*CarE*) (GSSPFG00010082001.3-RA, GSSPFG00020288001.5-RA, GSSPFG00000131001.3-RA), adipokinetic hormone (*AKH*) and adipokinetic hormone receptor (*AKHR*) in *Clo-rr S. frugiperda* is an additional mechanism to contribute to *S. frugiperda* resistance to chlorpyrifos. The enhanced activity of carboxylesterases was proved to be the mechanism of chlorpyrifos resistance in *N. lugens* (Tang et al., 2020a). Experiments for *AKH* and *AKHR* knockdown led to decreased CarE activity, and loss of resistance to chlorpyrifos in *N. lugens*. The role of AKH in regulating CarE availability and *N. lugens* resistance to chlorpyrifos was further demonstrated by AKH injections into susceptible *N. lugens* (Tang et al. 2020a). The activation of CarE expression and activity by AKH in insects exposed to chlorpyrifos leading to insecticide resistance was also demonstrated for *Bactrocera dorsalis* (Diptera: Tephritidae) resistant to another organophosphate (malathion) (Yang et al., 2021). AKH regulation of detoxifying enzymes was also detected for CYP enzymes, but in this case the inhibition of AKH upon exposure to the neonicotinoid imidacloprid induced the production of reactive oxygen species (ROS). ROS activated the transcription factors cap ‘n’ collar isoform-C (CncC) and muscle aponeurosis fibromatosis (Mafk), which in turn induced expression of CYP6ER1, while ROS directly mediates the activation of CYP6AY1 expression. Both CYP enzymes were shown to confer resistance to imidacloprid in *N. lugens* (Tang et al., 2020b).

AKH has also been shown to have a protective role in response to oxidative stress. AKH overexpression was shown to decrease the amount of protein carbonyls in hydrogen peroxide-challenged flies. Flies overexpressing AKH also had much higher levels of the major antioxidant protein sestrin when compared to AKH-RNAi flies. Sestrin is under regulation of Forkhead box class O transcription factor (FoxO), implying FoxO activation of sestrin occurs downstream of AKH to offer protection against increased oxidative stress (Bednářová et al., 2015). The overexpression of FoxO in *Clo-rr* strain associated with the positive regulation of FoxO by AKH, and their association with oxidative stress response indicates resistance of *Clo-rr* is also supported by an increased protective oxidative response, with the consequent inhibition of cell apoptosis (Yang et al., 2021).

The overexpression of G protein-coupled receptors (GPCRs) also support the role of FoxO as a mechanism of resistance of *S. frugiperda* to chlorpyrifos. GPCRs are important to environmental and physiological signaling throughout organismal life cycle, being activated by neuropeptides and peptide hormones (Hewes and Taghert, 2001; Park et al., 2002; Hansen et al., 2010). GPCRs are responsible to trigger a cascade of transduction by activating adenylate cyclase (ADCY), which in turn leads to increased cAMP synthesis and concentration in the cell (Hepler and Gilman, 1992). cAMP elevation directly affects the forkhead box class O (FoxO) transcription factor, resulting in the up-regulation of anti-oxidative enzyme genes discussed above (Bednářová et al., 2015; Kodrík et al., 2015).

In conclusion, the intensive use of lambda-cyhalotrin and chlorpyrifos in *S. frugiperda* control in Brazil led to an increased probability of survival in field populations, resulting in the selection of resistant strains. The molecular mechanism of resistance of *S. frugiperda* strains to both insecticides is polygenic, physiological in nature. Resistance of *S. frugiperda* to lambda-cyhalothrin occurs due to an elevation of the detoxification process by an increased expression of *CYP3* and *CYP6* genes. The mechanism of resistance to chlorpyrifos involves the up-regulation of several detoxification enzymes (CYP, UGTs and CarE), and of the protective oxidative response through up-regulation of AKH and FoxO. Our transcriptome analyses provided valuable information for the comprehension of the resistance mechanisms of *S. frugiperda* to lambda-cyhalothrin and chlorpyrifos. Such results will allow the development of management strategies to delay events of field-evolved resistance that, if implemented by regulatory agencies, the industry and growers will reduce non-target effects of pesticides, reduce the cost of crop production and reduce the risks to food security.

## Acknowledgements

We thank PROMIP (SISBIO License #40380-5) for helping to collect insect samples. Research was carried out using the computational resources of the Center for Mathematical Sciences Applied to Industry (CeMEAI) funded by FAPESP (grant 2013/07375-0).

## Contributions

ARBN, FLC, and CO conceived the study. ARBN, KLSB and JGR collected the data. ARBN performed the analysis. ARBN, FLC, RHK, and FSAA wrote the main manuscript. All authors contributed to writing and editing the manuscript.

## Data Availability Statement

Illumina data will be publicly available at the NCBI BioProject PRJNA780767.

## Ethical approval

This research did not involve humans or vertebrates.

